# Natural Variations of Spring Wheat Nitrogen and Carbon Assimilation under Different Inorganic Nitrogen Forms and CO2 levels

**DOI:** 10.64898/2026.04.28.721410

**Authors:** Pornpipat Kasemsap, Junli Zhang, Daniel J. Kliebenstein, Arnold J. Bloom

**Affiliations:** Department of Plant Sciences, University of California, Davis, CA 95616 USA; Texas A&M AgriLife Research, Texas A&M AgriLife High Plains Research and Extension Center, 3211 Russell Long Blvd, Canyon, TX 79015

**Keywords:** breeding, cereal, climate change, genes, GWAS, mapping, nitrogen, nitrogen use efficiency, wheat, yield

## Abstract

Wheat (*Triticum aestivum* L.) production depends upon nitrogen fertilization and may be threatened by the climate conditions anticipated in the next few decades. The natural variation in the ability of wheat to assimilate carbon and different inorganic nitrogen forms, nitrate (NO_3_^-^) and ammonium (NH_4_^+^), into vegetative growth remains unexplored. Here, we evaluated growth under either NO_3_^-^ or NH_4_^+^ as a sole nitrogen source for 875 spring hexaploid wheat accessions that represent the genetic diversity within the global germplasm. These accessions varied over 8-fold in vegetative biomass but grew similarly under moderate levels of either nitrogen form. At high, potentially toxic concentrations of NH_4_^+^, however, they lost approximately 20% of their biomass. We characterized the influence of changing CO_2_ levels in bi-parental Nested Association Mapping populations. Genetic backgrounds determined wheat biomass responses to CO_2_ enrichment and nitrogen form. Genome-Wide Association and linkage mapping identified certain loci as consistently associated with biomass accumulation under different nitrogen forms and CO_2_ levels. These results will assist breeding efforts to develop food crops that are resilient to the world’s climate changes.

## Introduction

Nitrogen, as an essential element for crop growth and food quality, is a major factor in global food security. Nitrogen use efficiency, however, the proportion of harvestable nitrogen in the final food product, remains below 50% (X. Zhang et al., 2021) and may decline under the climate conditions anticipated in the next few decades (Ren et al., 2023). Nitrogen fertilization itself is responsible for about 5% of the anthropogenic greenhouse gas emissions that drive global climate change (Gao & Cabrera Serrenho, 2023). Production of food crops like wheat (*Triticum aestivum* L.), the major plant source of protein and calories in the human diet, relies heavily on nitrogen fertilization (Hawkesford, 2014). Successful deployment of wheat cultivars with improved nitrogen use efficiency may simultaneously mitigate climate change and ensure global food security.

Plants convert the major soil inorganic nitrogen forms, ammonium (NH_4_^+^), or nitrate (NO_3_^-^), into organic forms such as proteins and nucleic acids necessary for life (Xu et al., 2012). Biomass production and nitrogen use efficiency are complex polygenic traits involving multiple genetic and physiological interactions that remain unresolved (Lammerts Van Bueren & Struik, 2017). To date, only a handful of genes that improve yield and nitrogen use efficiency have been incorporated into breeding programs; these involve vegetative tiller development, flowering time, and ammonium assimilation (Kasemsap & Bloom, 2022). Because up to 90% of grain nitrogen is relocated from growing tissues prior to anthesis (Kong et al., 2016), vegetative biomass accumulation is critical to both grain yield and protein concentration. We lack information about the genetic architecture of biomass production under different inorganic nitrogen sources, although such information would facilitate the breeding of crops with improved nitrogen use efficiency.

NH_4_^+^ nutrition may be advantageous over NO_3_^-^ nutrition for multiple reasons. First, plants require less energy to assimilate NH_4_^+^ than NO_3_^-^ into amino acids (Bloom, 2010; Cox & Reisenauer, 1973). Second, NH_4_^+^ binds to soil particles in temperate soils because they are usually negatively charged; in contrast, NO_3_^-^ remains mobile in the soil solution and may leach away from the root zone before being absorbed by a crop (J. Norton & Ouyang, 2019). Third, as a result of the lower soil mobility of NH_4_^+^, NH_4_^+^-based crop production systems emit less NO_2_, a highly potent greenhouse gas, to the atmosphere. Fourth, elite wheat varieties that were modified to sustain NH_4_^+^ in the root zone through biological nitrification inhibition yielded more grain without sacrificing grain quality (Subbarao et al., 2021). Lastly, wheat under the elevated atmospheric CO_2_ levels anticipated in the near future maintained grain protein and nutritional qualities when it received NH_4_^+^, but not when it received NO_3_ ^-^ (Rubio-Asensio & Bloom, 2016).

Nevertheless, plants generally cannot tolerate high concentrations of NH_4_^+^ (Britto & Kronzucker, 2002). Although NH_4_^+^-based fertilizers are widely used globally (Nishina et al., 2017), rapid microbial nitrification of NH_4_^+^ into NO_3_^-^ in aerobic agricultural soils typically results in NO_3_^-^being the prevalent form available to crops (J. M. Norton, 2015). Most previous genetic studies on nitrogen use efficiency primarily focused on varying nitrogen levels (Cormier et al., 2016). A few studies reported contrasting responses to different nitrogen forms based on results from limited genetic materials (Cox & Reisenauer, 1973; Fuertes-Mendizábal et al., 2013; C. O’Sullivan et al., 2016; Sun et al., 2013). The extent to which the current wheat germplasm utilizes either form of inorganic nitrogen or tolerates high NH_4_^+^ levels essentially remains unexplored.

Association mapping enables examination of the regulatory genetic elements underlying phenotypic changes. Genome-wide association (GWA) mapping overcomes the limited recombination events and genetic variation of a traditional linkage mapping (Korte & Farlow, 2013). From 2009 to 2020, wheat GWA studies cumulatively reported ∼47,000 putative loci for a wide range of traits (Saini et al., 2022). Yet GWA analysis alone remains insufficient to identify the causal loci (McCarthy & Hirschhorn, 2008), especially for traits with complex genetic architecture, even those with high heritability (Korte & Farlow, 2013). Additional high-resolution mappings can provide further validation and functional characterization. For example, combining the quantitative trait loci (QTL) information from a GWA panel with a linkage mapping improves the design of wheat breeding pipelines (Wang et al., 2023). Rather than deploying genome-wide mapping as a sole approach, successful endeavors initiated GWA as an exploratory tool followed by cross-validation with linkage mapping of different plant materials such as nested-association mapping (NAM) populations (J. Zhang et al., 2018). NAM populations (Blake et al., 2019; McMullen et al., 2009; Yu et al., 2008) provide a high mapping resolution from the recombinant inbred lines as well as rich allelic variations from multiple founder parents (Saini et al., 2022).

Here, we employed diverse genetic materials to characterize the genetic basis of wheat carbon and nitrogen assimilation under ammonium versus nitrate as a sole nitrogen source. First, we explored the vast diversity within the global germplasm and mapped loci associated with biomass production under ambient CO_2_ environments via a GWA analysis. Second, we used bi-parental nested association mapping populations to validate and narrow down candidate loci and to evaluate further how CO_2_ enrichment alters responses to different nitrogen sources. Our study established the extent to which wheat utilizes ammonium or nitrate as a major nitrogen source, including potentially toxic concentrations of NH_4_^+^. We identified candidate loci that can guide breeding food crops that are resilient to future climate conditions.

## Materials and Methods

### National Small Grains Core Collection at ambient CO_2_

We grew a panel of 875 spring hexaploid wheat (*Triticum aestivum* L.) accessions from the USDA-ARS National Small Grains Core Collection (NSGCC) that included 154 landraces, 261 registered cultivars, and 460 actively cultivated materials, which represent the germplasm in 89 countries worldwide (Maccaferri et al., 2015). We disinfected mature seeds collected from field and greenhouse experiments in Davis, CA with 10% v/v NaHClO_3_ for 10 minutes, rinsed them multiple times with deionized water, and placed them on a germination paper soaked with deionized water. We kept rolls of the germination paper at 4°C for at least 48 hr and later moved them to room temperature for 4 d until transplanting. We transplanted seedlings into a 20 dm^3^ tub filled with a modified Hoagland solution (Epstein & Bloom, 2005) containing either NH_4_^+^ (0.5 mM or 20 mM) or NO_3_^-^ (0.5 mM) as a sole nitrogen source. The solution pH was initially adjusted to 6.0, and the solution was replenished twice in the first week, and every 1 to 2 days during the second week of growth. We grew seedlings at ambient CO_2_ concentrations in an evaporative cooled, ventilated greenhouse at University of California, Davis (38.5365 N, 121.7470 E) and harvested the whole plant at the tillering stage (Zadoks 21-25 or Feekes 2-3) 15 d after transplanting, when growth differences due to nutrient treatments became clearly visible.

Our experiment consists of 6 replicates: 3 replicates grown consecutively during summer months from June to August of 2016 and again in 2017. In each replication, we randomly divided the whole panel into 5 groups; each group was germinated and transplanted into the same greenhouse consecutively one day after the other for 5 sequential days. We retrieved weather records from a local CIMIS station (38.5358 N, 121.7764 E) located 2.55 km from the greenhouse (https://cimis.water.ca.gov/). Seasonal solar radiation levels and temperatures characterized each summer month (June 2016, July 2016, August 2016, June 2017, July 2017, and August 2017) as a distinct environment, resulting in 6 growth environments.

### Genome-Wide Association mapping

We calculated BLUP (Best linear unbiased prediction) for each genotype in a mixed linear model using the function lmer in R/lme4 (Bates et al., 2015) where we specified year, replicate, germination group, and blocks within each replicate as random effects. The mixed linear model was as followed for each nitrogen treatment: mass ∼ genotype × nitrogen + (1|year) + (1|year: replication) + (1|year: replication: germination group) + (1|year: replication: germination group: block). We then used the fitted biomass under each nutrient treatment, corrected by the biomass of the reference genotype Veery-10 in each germination group, to calculate the dry mass ratio of interest. To determine a broad sense heritability and the proportion of variance explained by each component, we ran a linear model with all factors as fixed effects and calculated type II sums of squares (Langsrud, 2003) using R/car version 3.1 (Fox & Weisberg, 2019) from an analysis of variance (ANOVA).

As inputs for Genome Wide Association (GWA) mapping with 4,585 SNP markers (Maccaferri et al., 2015), we used the mean of total plant dry mass under NH_4_^+^ and NO_3_^-^, and the ratio of the dry mass of plants under different concentrations and forms of nitrogen, across 3 replicates in each year and 6 replicates over the 2 years. We performed GWA of each biomass trait with the K (kinship matrix) in a mixed linear model (MLM) in R/GAPIT (Lipka et al., 2012), as described previously to be most the suitable for this mapping population (Maccaferri et al., 2015). MLM corrected with population structure (Q) and principal components (10), as well as BLINK (Huang et al., 2019) without any correction besides K resulted in generally the same set of significant QTL (data not shown) under our stringent threshold. For each trait, we identified SNPs that were significant (*P* < 0.05, marker-wise) in at least one replicates in any year and with at least one replicate, or the means across replicates or years, being highly significant overall (*P* < 9.65 x 10^-5^, experiment-wise), given α = 0.1 for the Bonferoni correction accounting for the number of tag-SNPs (1036 SNPs). The 50% LD decay rate is 1 cM in this diversity panel when genotyped with the 9K SNP array because of the rich diversity and high level of historical recombination (Maccaferri et al., 2015). We set the confidence intervals for the QTL-harboring regions at ± 1.6 cM, corresponding to the critical level of r^2^ = 0.3 (Maccaferri et al., 2015).

### Bi-parental nested association mapping populations at ambient CO_2_

We grew eight biparental NAM populations, each including 75 recombinant inbred lines (RILs), to validate the significant SNPs detected in the GWA. These populations were generated from crosses between the CIMMYT line Berkut (Irene/Babax//Pastor) as a female parent and eight lines (PBW343, Dharwar Dry, LR23 (PI 70613), LR3 (CItr 7635), RAC875, Kern, RSI5, and UC1419 (Patwin-515HP or PVP 201600390)) with diverse genetic backgrounds as male parents (Blake et al., 2019). These lines were genotyped with the Illumina wheat iSelect 90K SNP array (Blake et al., 2019; J. Zhang et al., 2018). We germinated and grew seedlings from July to September 2020 in the same greenhouse environments as the GWAS experiments described above. In each tub of the 25 genotypes being evaluated, we added reference genotypes: 3-5 plants of cv. Veery-10 and 2-3 plants of cv. Berkut and Hahn 1RS.

### Linkage and joint linkage-association mapping

We calculated coefficients of dry mass for each genotype in a mixed linear model using the function lmer in R/lme4 (Bates et al., 2015) where we specified replicate and blocks within each replicate as random effects, genotype as a fixed effect, and only used the mean biomass of cv. Veery-10 in each tub as a covariate to correct for dry mass variation across blocks in different parts of the greenhouse. The mixed linear model for each nutrient treatment was mass (mg) ∼ dry mass of Veery + genotype + (1|replication) + (1|replication:block). We then used the fitted biomass in each nutrient condition to calculate the dry mass ratio of interest.

As input for linkage mappings, we used an individual replication value, mean dry mass under NH_4_^+^ and NO_3_^-^, and dry mass ratio of plants under different concentrations and forms of nitrogen, of the 3 repeated experiments across summer months, each with 1 plant replicate. We performed QTL mapping for each trait in each individual population in R/qtl (Arends et al., 2010) and joint linkage-association mapping for all 8 populations in TASSEL, using Stepwise regression association analysis (Bradbury et al., 2007). For each trait, we identified SNPs with LOD > 2.0 in each population, or with p-value < 0.001 in joint linkage-association mapping.

### Bi-parental nested association mapping populations at elevated CO_2_

To evaluate how biomass changes under CO_2_ enrichment, we germinated seedlings of six bi-parental nested association mapping populations with the same protocol above, but transplanted them into nutrient solution tubs placed in controlled environment facilities. The growth conditions in controlled environmental chambers (Conviron, Winnipeg, Canada), equipped with non-dispersive infrared analyzers, were as follows: 750 ppm CO_2_ (concentration expected at the end of the 21^st^ century (IPCC, 2021)), 50% relative humidity, 16 hr of 800 µmol m^-2^ s^-1^ PPFD at canopy height with 22/15°C light/dark temperature. We evaluated 25 genotypes in each tube, including 4 plants of the reference genotype cv. Veery-10.

We calculated coefficients of biomass traits with the same approach as described above: we specified replicate and blocks within each replicate as random effects, genotype as a fixed effect, and used the mean biomass of Veery-10 in each tub as a covariate to correct for dry mass variation across replications. The mixed linear model was as follows for each nutrient treatment: dry mass (mg) ∼ dry mass of Veery + genotype + (1|replication) + (1|replication:block). We then used the fitted biomass in each nutrient condition to calculate dry mass ratio of interest. We performed QTL and joint linkage with the same approach as described above. We compared the significant QTL between ambient and elevated CO_2_ levels.

### Data processing and visualization

Unless otherwise noted, we performed all analyses in R (R Core Team, 2023) in an integrated development environment Rstudio (RStudio Team, 2020). We processed data with R/tidyverse version 2.0 (Wickham et al., 2019) and R/data.table version 1.14.8 (Dowle & Srinivasan, 2023), and visualized most of the numerical data with R/ggplot2 version 3.4.4 (Wickham, 2016). We employed packages R/see version 0.8.1 (Lüdecke et al., 2021), R/ggbeeswarm version 0.7.2 (Clarke et al., 2023), R/corrplot version 0.92 (Wei & Simko, 2021), and R/rMVP version 1.0.6 (Yin et al., 2020) for half violin plots, scattered violin plots, correlation plots, and Manhattan plots respectively.

## Results

### Wheat tolerated high levels of NH_4_^+^and adapted to using either nitrogen forms

Whole seedling biomass of the worldwide spring wheat collection in an ambient CO_2_ environment spanned 8-fold from 0.1 g to over 0.8 g (Figure 1). Monthly differences (experimental replicate) primarily accounted for the greatest portion of this biomass variation (Figure 1B). Biomass declined (Figure 1C) as light availability declined (Figure 1A), suggesting a response to changes in day length and temperature from the beginning of the summer (June) to late in the season (August) across both years of the experiment (Figure 1C).

**Figure 1.**
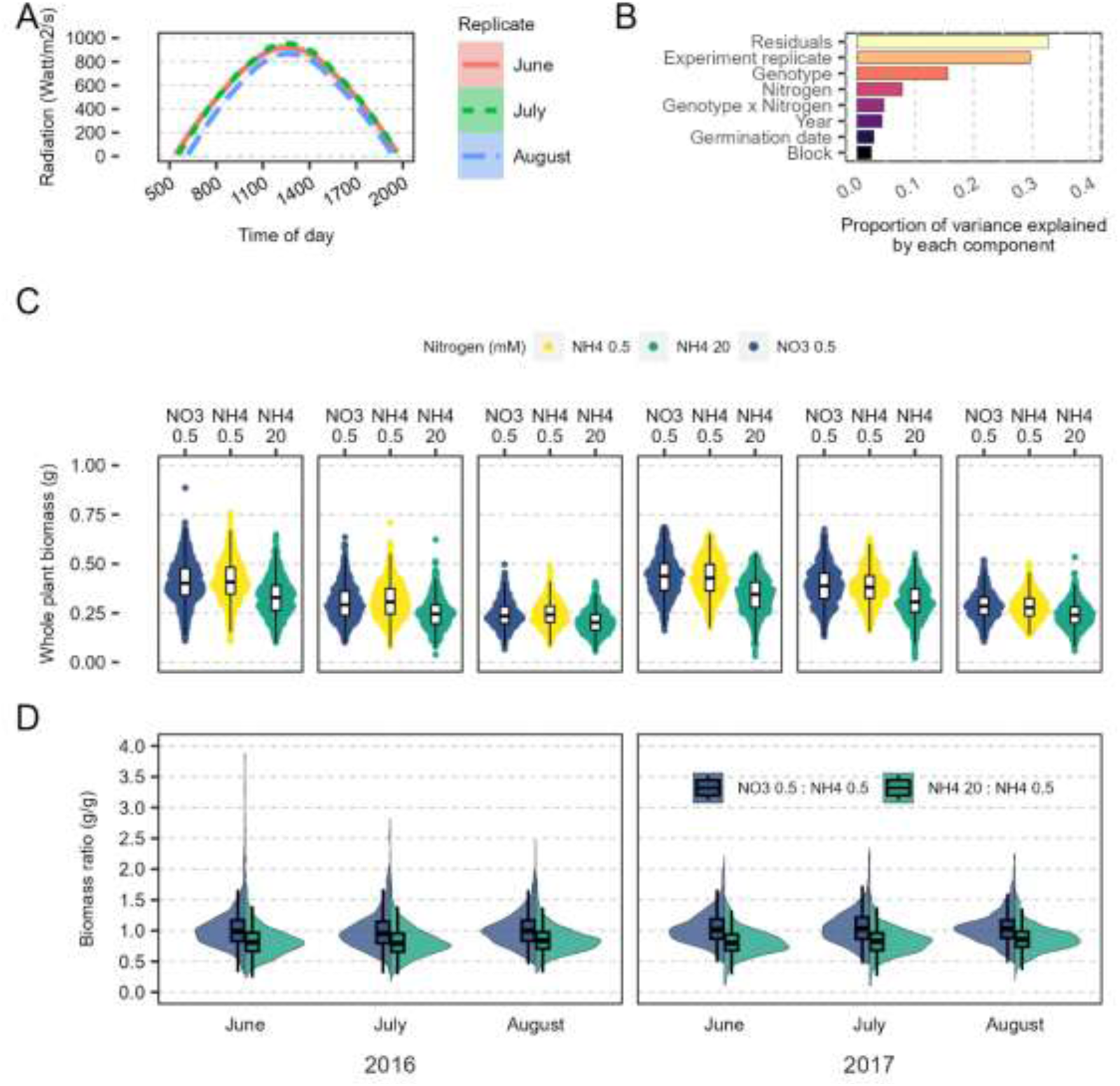
Wheat biomass varied with seasonal changes but nitrogen responsiveness to either form of nitrogen and high NH_4_^+^ level remained consistent. (A) Seasonal variations in radiation, averaged across 2016 and 2017 (Watt m^-2^ s^-1^), (B) Proportion of variance explained by each component in the experiment, (C) Distribution of whole plant biomass of 875 global wheat accessions receiving NO_3_^-^ (0.5 mM) or NH**_4_^+^** (0.5 or 20 mM) as a sole nitrogen source at 15 d after transplanting under ambient CO_2_ concentrations) over 6 independent replicates in Summer months of 2016 and 2017. The box plots represent minima, 25^th^ percentiles, medians, 75^th^ percentiles and maxima n=875 genotypes with 1 plant replicate in each month and year combination). Each dot represents a plant replicate. (D) Distribution of whole plant biomass ratio of 875 global wheat accessions under NO_3_^-^ (0.5 mM) or NH**_4_^+^** (20 mM) to the reference treatment NH**_4_^+^** (0.5 mM) over 6 independent replicates in summer months of 2016 and 2017. The box plots represent minima, 25^th^ percentiles, medians, 75^th^ percentiles and maxima n=875 genotypes with 1 plant replicate in each month and year combination). Figure C and D share the same x-axis.

The response to nitrogen form was consistent across seasonal changes (Figure 1D). We defined a moderate level 0.5 mM NH_4_^+^ as a reference treatment and compared the responses between nitrogen conditions by calculating the biomass ratio of the same genotype to the reference treatment. The biomass ratio under 0.5 mM NO_3_^-^ to that under 0.5 mM NH_4_^+^ averaged about 1.0; the responses ranged from ∼0.25 to ∼1.5, suggesting potential adaptations to the specific nitrogen forms. Under 20 mM NH_4_^+^, the average response was about 80% of the biomass under 0.5 mM NH_4_^+^ and ranged from about 0.1 to 1.5, indicating larger variation in tolerance to high NH_4_^+^ levels within the germplasm. Seven subgroups (Maccaferri et al., 2015) within the germplasm that was used to correct for the population structure displayed slight different to nitrogen responses, but there is no clear association with the geographical origins of the accessions (Supplementary Figure 1).

### Biomass is controlled by multiple loci across different nitrogen conditions

We performed genome-wide association analyses with biomass of each individual experiment replicate and across the whole experiment, accounting for biomass plasticity across the season. In total, 2419 SNPs were significantly associated with biomass in any given nitrogen condition and environment at the marker-wise threshold (p<0.05) (Supplementary Table 1). To narrow down candidate loci and limit false positive, we considered SNPs that were highly significant in at least one environment under a stringent experiment-wise p-value threshold (∼p<0.0001). Experiment-wise, we identified 20 significant SNPs representing 14 loci that were either unique to or shared between the nitrogen conditions across multiple environments (Table 1). Different sets of SNPs were associated with biomass and biomass ratio. The majority of the significant SNPs were identified under 20 mM NH_4_^+^. Two loci associated with SNPs IWA2077 and IWA6647 were significant for biomass in all nitrogen treatments.

**Table 1.**
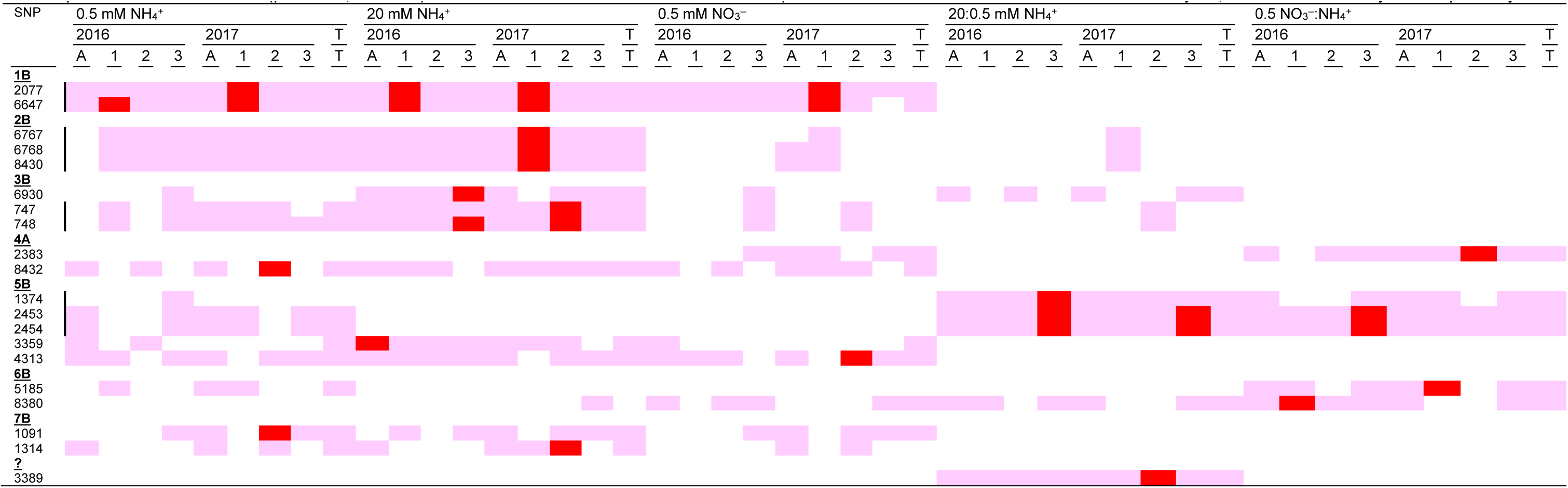
Significant SNPs consistently associated with dry mass and dry mass ratio of wheat receiving 0.5 mM NH4^+^, 20 mM NH4^+^ or 0.5 mM NO3^-^ at the marker-wise threshold (p<0.05; light red cells) and experimental-wise threshold (p<0.0001; red cells). Year and environment are indicated on the top rows. A and T indicates overall means within each year, and across the two years respectively.

### Biomass partitioning varied upon nitrogen sources, CO_2_ levels and genetic backgrounds

The bi-parental populations, however, only presented a portion of the phenotypic variation previously observed in the global diversity panel (Figure 1). Genetic backgrounds and nitrogen conditions influenced the biomass of these bi-parental populations (Figure 2). Like the GWAS panel, season variation across multiple independent replicates significantly influenced the overall biomass (Figure 2A-B). Population Berkut x RAC875 produced the highest amount of biomass (Figure 2C). At the whole plant level, all populations exhibited comparable responses to nitrogen treatments (Figure 2C). Nitrogen responsiveness, however, varied between plant organs (Figure 2B). Interestingly, NO_3_^-^-grown plants generally maintained comparable root growth to NH_4_^+–^-grown plants, but at a cost of the shoot growth (Figure 2E-F). As a result, seedlings under NO_3_^-^ had lower whole plant biomass, but still partitioned more biomass into root than plants under NH_4_^+^ (Figure 2D,G). In contrast, biomass partitioning under different levels of NH_4_^+^ remained comparable for most populations (Figure 2G); the lower whole plant biomass under the high NH_4_^+^ was due to slower growth of both shoots and roots (Figure 2E-F).

**Figure 2.**
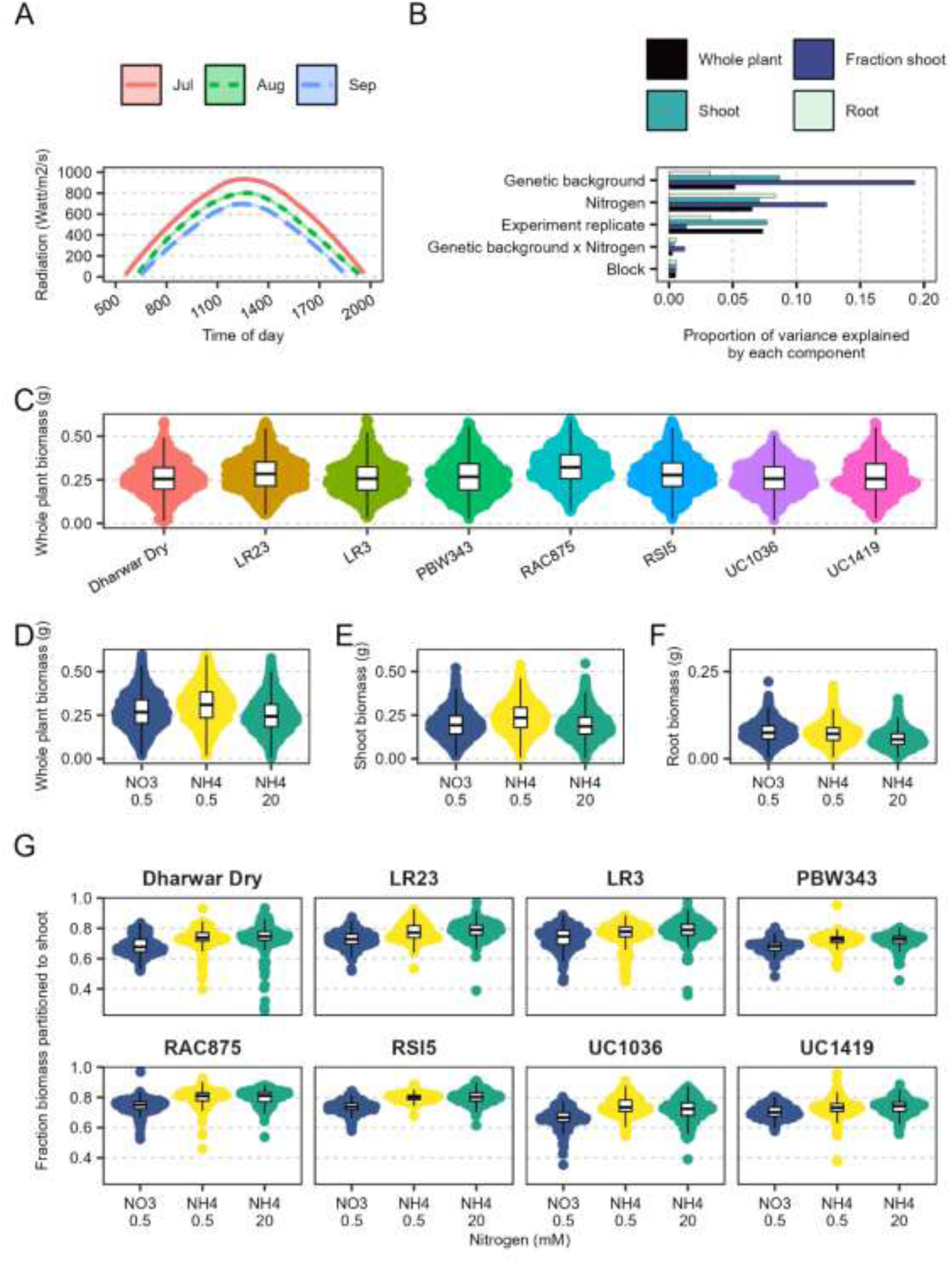
Genetic backgrounds determined biomass partitioning under different nitrogen source. (A) Seasonal radiation variations in 2020 (Watt m^-2^ s^-1^), (B) Proportion of variance explained by each component in the experiment. Each box represents different plant parts, including whole plant, shoot, root, and fraction of biomass partitioned to shoot. Variances explained by residuals are not shown. The sum of all components within the same plant part is 1. (C-G) Distribution of biomass by (C) population, (D-F) plant parts by nitrogen conditions (whole, D; shoot, E; root, F), and (G) fraction of biomass partitioned to shoot by nitrogen conditions of 8 bi-parental populations receiving NO_3_^-^ (0.5 mM) or NH**_4_^+^** (0.5 or 20 mM) as a sole nitrogen source at 15 d after transplanting under ambient CO_2_ concentrations over 3 independent replicates in summer months of 2020. The box plots represent minima, 25^th^ percentiles, medians, 75^th^ percentiles and maxima. Each dot represents an average of an individual genotype. (n=75 genotypes with up to 3 replicates). Although biomass significantly varied across environments as illustrated in (B), we showed here in C-G aggregated values across the environments. For QTL mapping, we used individual values for each environment to account for response plasticity across the season.

Comparing growth of the bi-parental populations under the ambient and elevated CO_2_ levels, the growth environments explained the largest variations (64%) in biomass (Figure 3A and Supplementary Figure 2). In the controlled environment facilities, several growth factors differed from the ambient conditions in the greenhouse, including higher light intensity and longer day length. Nitrogen source and its interaction with CO_2_ further determined the biomass responses to CO_2_ enrichment (Figure 3A-B). Under ambient CO_2_, the biparental populations were relatively more tolerant to high NH_4_^+^ and grew faster under NH_4_^+^ than under NO_3_^-^. In contrast, NO_3_^-^ nutrition boosted higher whole plant biomass under CO_2_ enrichment, showing a different nitrogen response than under the ambient CO_2_ condition. The high NH_4_^+^ level also slowed down growth under CO_2_ enrichment for all populations, except for population PBW343.

**Figure 3.**
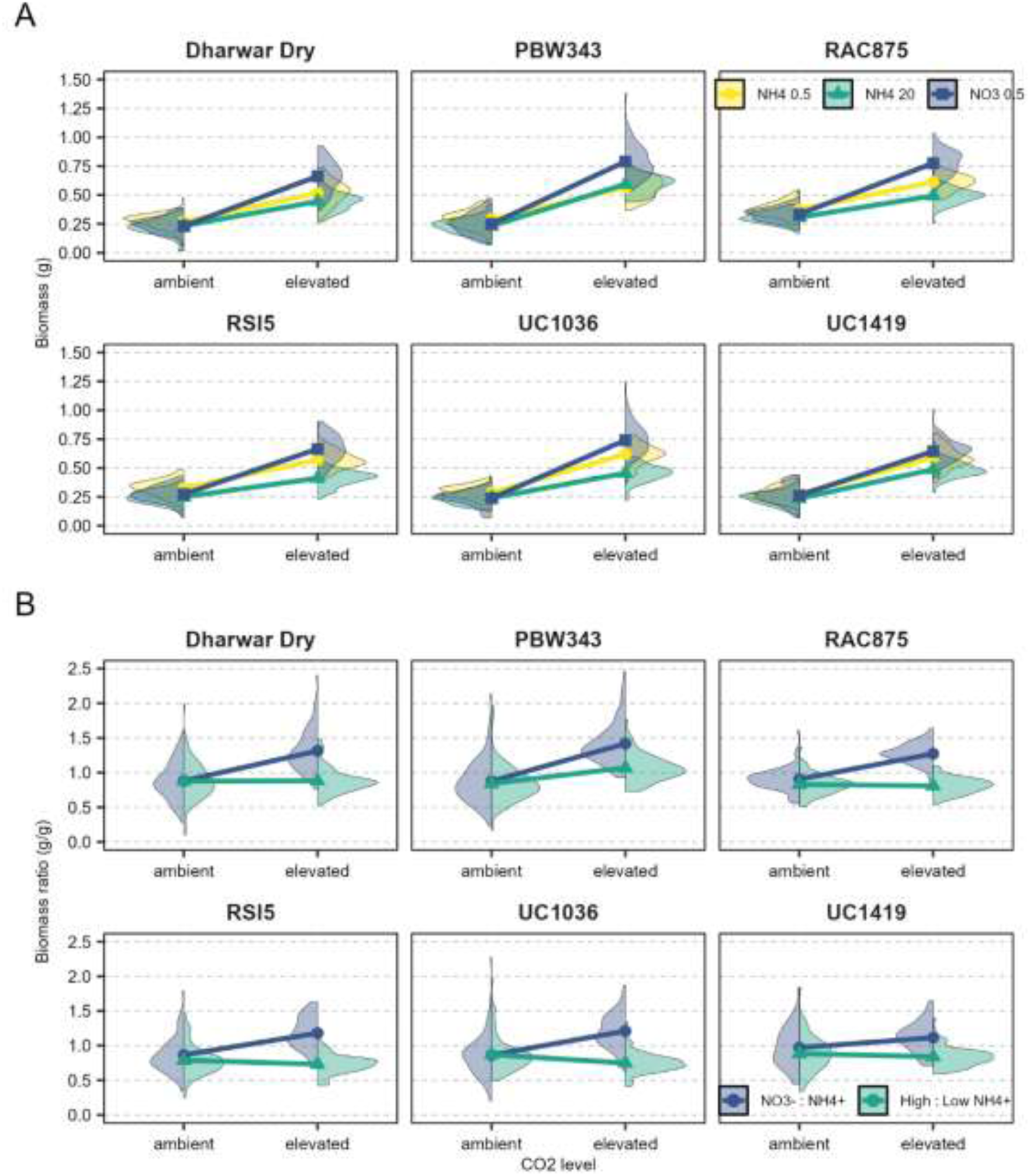
CO2 enrichment changed wheat responses to inorganic nitrogen forms. (A) Distribution of whole plant biomass of 6 bi-parental populations receiving NO_3_^-^ (0.5 mM) or NH**_4_^+^** (0.5 or 20 mM) as a sole nitrogen source at 15 d after transplanting under ambient and 750 ppm CO_2_ (n=75 genotypes with 4 plant replicate for each population). (B) Distribution of whole plant biomass ratio of 6 bi-parental populations under NO_3_^-^ (0.5 mM) or NH**_4_^+^** (20 mM) to the reference treatment NH**_4_^+^** (0.5 mM) (n=75 genotypes with 4 plant replicate for each population). Symbols represent the average responses under different nitrogen conditions. Lines connect the averages between the two CO_2_ levels.

Correlations between biomass traits of NAM populations suggest genetic-dependent changes in carbon and nitrogen assimilation across the two CO_2_ levels (Supplementary Figure 3 and Supplementary Figure 4). Biomass across different nitrogen conditions were positively correlated, with coefficients >0.78. NO_3_^-^ preference, defined as the biomass ratio of growth at 0.5 mM NO_3_^-^ over the growth at 0.5 mM NH_4_^+^, was also positively correlated with NH_4_^+^ tolerance, defined as the biomass ratio under high and moderate NH_4_^+^ concentrations (Supplementary Figure 3A). For individual populations, the correlations of biomass were also positive across nitrogen conditions (Supplementary Figure 3B-G). The correlations between biomass ratio were consistent in direction but varied in magnitude. NH_4_^+^ tolerance was generally not correlated with growth under NO_3_^-^, except for population UC1036 (Supplementary Figure 3F) and population PBW343 (Supplementary Figure 3C).

### Linkage mapping further identified new loci that are unique to genetic backgrounds

Comparison between bi-parental populations further suggested that responses to different nitrogen forms may be primarily controlled by distinct sets of genes specific to genetic backgrounds. We identified 301 unique significant markers from the linkage mapping of individual populations, although the joint linkage mapping did not detect any common QTL that were shared across all bi-parental populations. We found only a few markers that were significant in multiple populations, although several markers were in close proximity to one another (Supplementary Figure 5). We detected 5 times more significant associations under the ambient CO_2_ level because we included biomass data from both shoot and root and from all individual experiment replicates across the season, as opposed to only the average across four replicates with similar controlled environment settings (Supplementary Figure 6). For further studies, we selected a subset of 121 QTL that had at least one significant marker with LOD > 2.0 (Supplementary Table 2). Among these markers, the additive effect sizes ranged widely from-32 to 61. The proportions of variance explained by the markers (R^2^) ranged from 0.12 to 0.21.

## Discussion

Plants optimize assimilation and allocation of resources to maximize growth (Bloom et al., 1985). During the Green Revolution, the success of modern wheat cultivars came at an expense of superfluous nitrogen fertilization (Li et al., 2018). Despite significant increases in harvest index and grain yield, the introduction of dwarf phenotypes into modern varieties also made crops less sensitive to changes in nitrogen inputs (Gooding et al., 2012; Li et al., 2018). The increase in atmospheric CO_2_ levels anticipated in the near future inhibits assimilation of NO_3_^-^ into protein (Bloom et al., 2010), but promotes carbon assimilation (Walker et al., 2021), thereby further exacerbating the imbalance of crop carbon and nitrogen. Here, we demonstrated how changes in rhizosphere nitrogen form influence shoot carbon assimilation into biomass, and how an increase in carbon supply of the magnitude anticipated by the end of this century alters responses to nitrogen. Combining multiple mapping populations, we identified and cross-validated candidate loci underlying wheat carbon and nitrogen assimilation.

Understanding the natural variation in a crop nitrogen utilization is fundamental for breeding for enhanced nitrogen use efficiency. We particularly highlighted the importance of genetic regulation of carbon-nitrogen relations. Here, we established a wide range of biomass responses to NO_3_^-^ versus NH_4_^+^ as nitrogen sources available within the global wheat germplasm. The available genetic resources offer great potential for improving nitrogen use efficiency to meet the challenge of the changing climate.

Interestingly, the bi-parental populations only represented partial genetic diversity available in the global wheat germplasm and were more adapted to using ammonium under ambient CO_2_ levels (Figure 2). Depending upon genetic background, the biomass cost of using a different nitrogen form may be as high from the effects of NH_4_^+^ toxicity, (Figure 2). Furthermore, genetic background also influenced responses to CO_2_ enrichment. At 750 ppm CO_2_, some genetic backgrounds grew even faster under an NH_4_^+^ concentration that was toxic to other genotypes (Figure 3). Because previous studies usually examined only a few genotypes, responses to nitrogen forms varied significantly (Cox & Reisenauer, 1973; Fuertes-Mendizábal et al., 2013; C. O’Sullivan et al., 2016; Sun et al., 2013). Employing a genetically diverse population, our study characterized both the general and the extreme responses to different nitrogen forms and NH_4_^+^ toxicity (Figure 1, Figure 2, and Figure 3). Genotypes with responses deviating from the averages may offer additional breeding values; for example, genotypes that tolerate the detrimental effects of elevated CO_2_ levels (Cassan et al., 2023). Our results emphasize that understanding nitrogen responsiveness will require examination of genetically diverse materials (Katz et al. 2022).

Here, we demonstrated that the majority of the global bread wheat germplasm employ both nitrogen forms effectively and maintained robust growth despite experiencing NH_4_^+^ toxicity (Figure 1). But, how should breeders make selection based on nitrogen form and NH_4_^+^ toxicity as a breeding target? The positive correlations of biomass under different nitrogen conditions support that prioritizing selection of high biomass, perhaps under ample nitrogen supply, should ensure sufficient nitrogen responsiveness across different soil nitrogen availability (Cormier et al., 2013). Here, NO_3_^-^ preference, defined as the ratio of biomass under both nitrogen forms at 0.5 mM, was also positively correlated with the biomass under 20 mM NH_4_^+^ for most populations (Supplementary Figure 3). Because these two values are independent, such relationship suggests connections between NO_3_^-^ preference and NH_4_^+^ tolerance. Even plants that enjoy NO_3_^-^ should generally perform decently under 20 mM NH_4_^+^. Such insights should allow breeders and farmers to maximize biomass production through manipulating soil nitrogen responses.

Breeding efforts guided by molecular genetic insights often attempt to alter expression of a few genes with large effects (Kasemsap & Bloom, 2022). Here, we demonstrated that small additive effects from a large number of regulatory elements may underlie biomass-nitrogen responsiveness. Heritability of wheat biomass is typically lower than other traits such as height or flowering time (Cormier et al., 2014). Still, the broad-sense heritability here is consistent with a previous field study determined at the same developmental stage (Molero et al., 2019). We identified large sets of candidate loci that were either unique to or stable across environments and mapping populations. Previous mappings in diversity panels of rice (Kasemsap et al., 2024) and Arabidopsis (Katz et al., 2022) underscore the complex genetics of nitrogen responsiveness. Relying on just a handful of genes may undermine the natural variations (Kasemsap & Bloom, 2022). A holistic understanding of the regulatory network that moderate carbon and nitrogen assimilation may better guide breeding for improved yield and grain quality simultaneously.

We identified both sets of candidate loci shared across and unique to nitrogen conditions across environments. Controlling false positive associations by enforcing stringent P-value thresholds may come at a cost of lower statistical power (Cormier et al., 2014). Here, we opted for a more lenient starting threshold with the Bonferroni correction, and prioritized associations that were consistent across multiple environments, which together were proven very stringent, yet effective in previous studies (Maccaferri et al., 2015; J. Zhang et al., 2018). Cross-validation between different mapping populations and studies are required to confirm the candidate loci identified in this study.

How adaptations to nitrogen form, NO_3_^-^ versus NH_4_^+^, influence nitrogen responsiveness remains inconclusive. Superior growth under specific forms perhaps reflects the genetic adaptation to resources in natural habitats. Wheat is often assumed to be more adapted to NO_3_^-^ perhaps because of the limited number of genotypes previously evaluated (He et al., 2022; S. Liu et al., 2019). In our study, the majority of the global germplasm perform well with either form of nitrogen (Figure 1), but the wide range of biomass responses in diverse genetic backgrounds also suggest potential adaptations (Figure 1 and Figure 2). For example, the natural abundance of rice *TCP19* functional allele follows gradients in soil nitrogen content and its gene expression follows changes in NO_3_^-^ levels (Y. Liu et al., 2021). Here, however, we did not observe any clear evidence for potential local adaptation in this diversity panel, despite its diverse geographic origins (Maccaferri et al., 2015). A growing body of evidence suggest that plants may regulate soil nitrification rates, thereby balancing the two inorganic forms to meet changing demands (He et al., 2022; C. A. O’Sullivan et al., 2016). For example, weed species (O’Sullivan et al., 2017) and wheat wild relatives (C. A. O’Sullivan et al., 2016) exude inhibitors to slow down the conversion of NO_3_^-^ into NH_4_^+^. Aligning crop nitrogen preference with fertilizer inputs and environments should enhance nitrogen use efficiency (S. Liu et al., 2019).

A myriad of interacting physiological and environmental cues influence differential responses to nitrogen source (Britto & Kronzucker, 2013). Here, the growing environments accounted for the largest proportion of biomass variations (Figure 1 and Figure 2). We also observed little to no significant correlation of biomass traits between growth in the greenhouse under ambient CO_2_ and in controlled environments under elevated CO_2_ (Supplementary Figure 4). Biomass values were correlated across nitrogen conditions only within the same growth conditions. Such relationships confirmed that nitrogen responsiveness is highly dependent on other environmental factors. Inclusion of the climate conditions anticipated in the near future, like elevated CO_2_ levels (Figure 3), is critical to assessing responsiveness to the inorganic nitrogen forms (Rubio-Asensio & Bloom, 2016). The influence of genetic adaptations to nitrogen sources on crop responses to climate change has significant implications on the future of food production and warrants further investigation.

## Acknowledgements

We appreciate the assistance during data collection of Joshua J. Claxton, Dana R. Lawrence, Kungyao Wang, Jordan A. Stefani, Carly M. Miranda, Elias Potashov, Guilherme M. Silva, Jenna O’Kelley, Sydney Koebel, James Schmidt, Anna M. Gomes, Ariel Herrera, Steven Moreno, Temiloluwa Salako, Zion Congrave-Wilson, Lisa Tam, and Khine Z. Lin. We thank Ron Lane, Bill Werner, Jiangao Chen, Davith Hin, and Antonio Lopez for their continued support at the greenhouse facilities and, Kevin Roberts and P. Andre Nelson for their continued support at the controlled environmental facilities. We thank Jorge Dubcovsky for the feedback on the project and the manuscript, Matthew E. Gilbert for discussion on the experimental set-up, and Daniel E. Runcie for discussion on the experimental set-up and statistical analyses for GWAS.

## Author contributions

Pornpipat Kasemsap and Arnold J. Bloom planned the research. Pornpipat Kasemsap collected data, analyzed data, and drafted the manuscript. Junli Zhang contributed to the genetic mapping. Daniel J. Kliebenstein contributed to statistical analyses and interpretation. All authors edited the manuscript.

## Data availability

The raw data that support the findings of this study will be deposited and openly available on Dryad. Analysis R codes are available at https://www.github.com/paulkasemsap/Wheat_Nform_Mapping.

## Supplementary Data

**Supplementary Figure 1.**
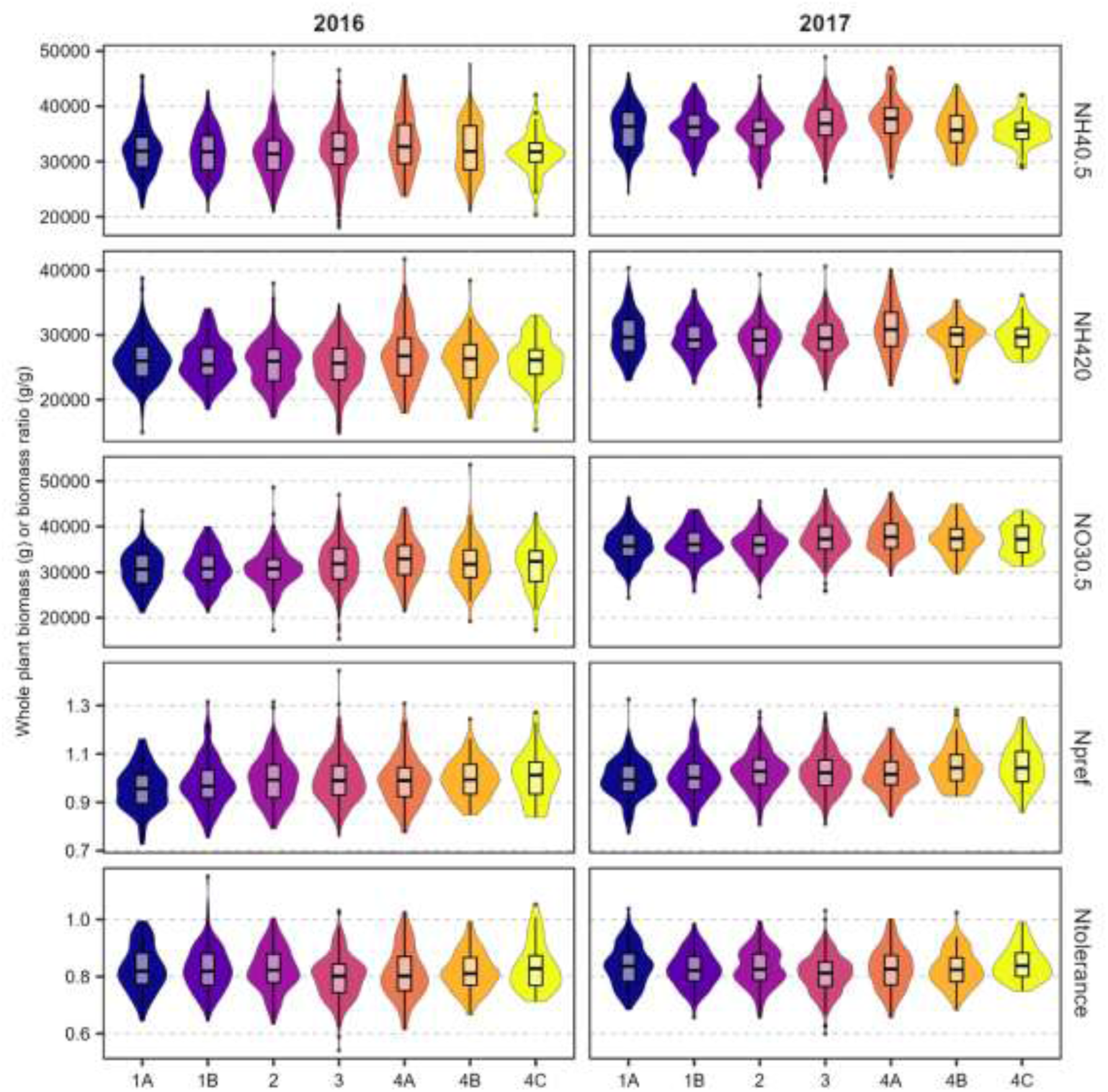
Distribution of biomass or biomass ratio by seven subpopulations within the diversity panel. The seven populations were identified in the analysis of the population structure of 875 accessions from the National Small Grains Collection spring wheat core collection. The accessions distributed across 87 countries on six continents. Subpopulation 1A is prevalent in Europe and Africa. Subpopulation 1B and 3 are prevalent in Europe. Subpopulation 2 is prevalent in America. Subpopulation 4A, 4B and 4C are prevalent in Asia. See Figure 1 and 2 (Maccaferri et al., 2015) for more details.

**Supplementary Figure 2.**
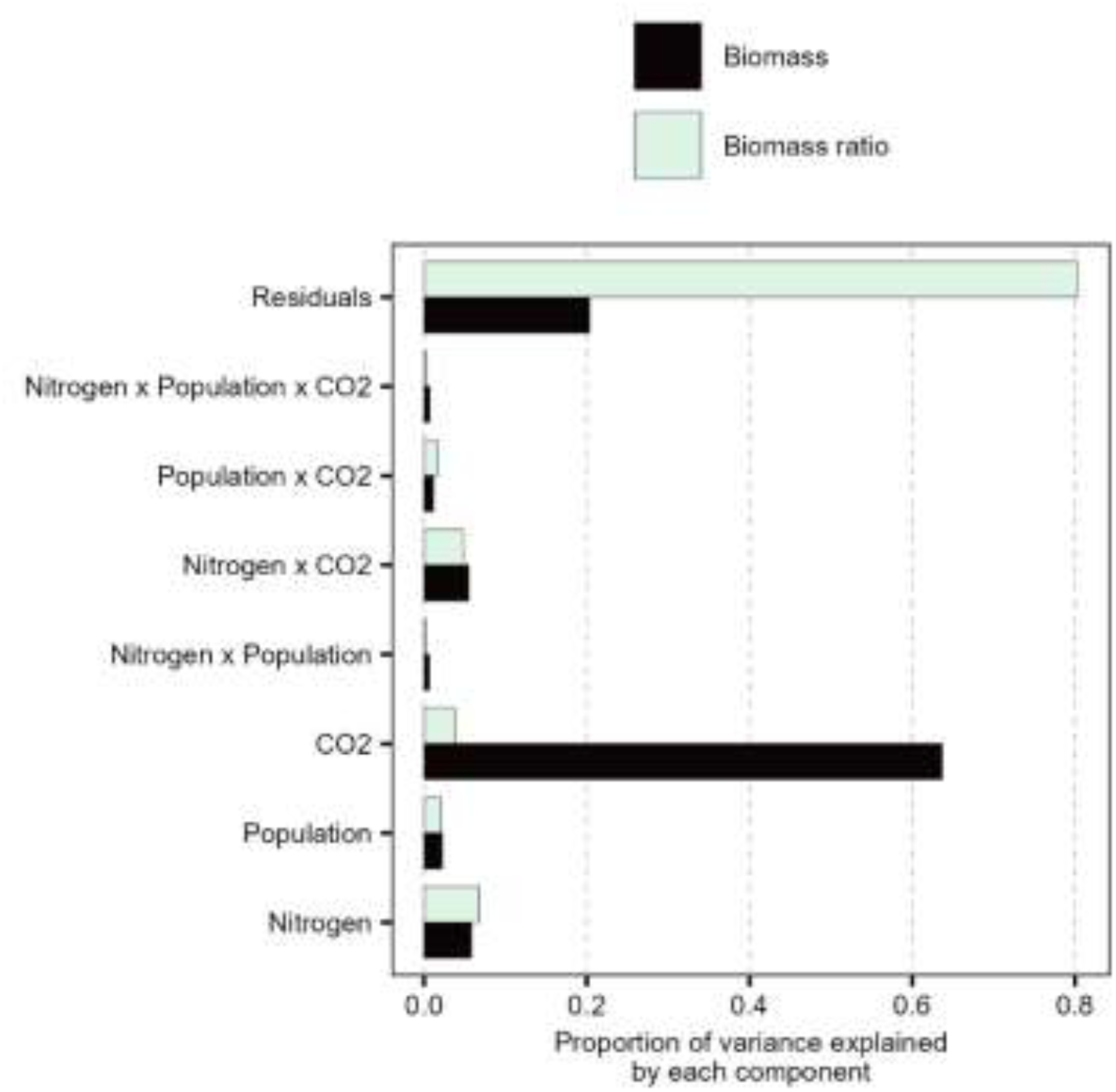
**Proportion of variance explained by each component in the Nested Association Mapping populations**

**Supplementary Figure 3.**
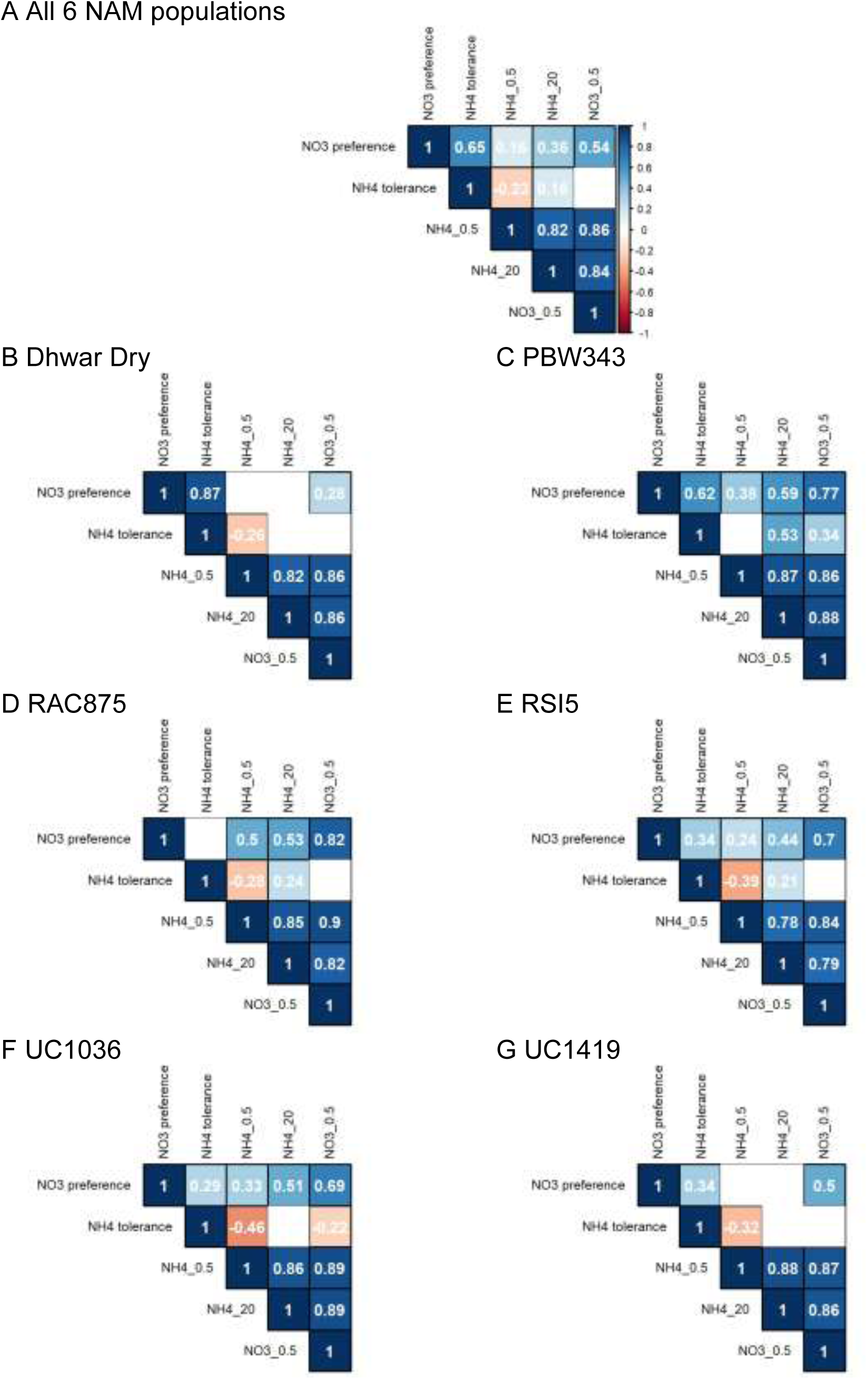
Correlation between biomass traits of 6 Nested Association Mapping populations. Pearson’s correlation coefficients at p-value < 0.01 between biomass traits across the CO_2_ concentrations (A) across 6 NAM populations and (B) – (G) by each NAM population. Insignificant correlations are not displayed. The horizontal bar displays reference color for negative correlations (red) to positive correlations (blue).

**Supplementary Figure 4.**
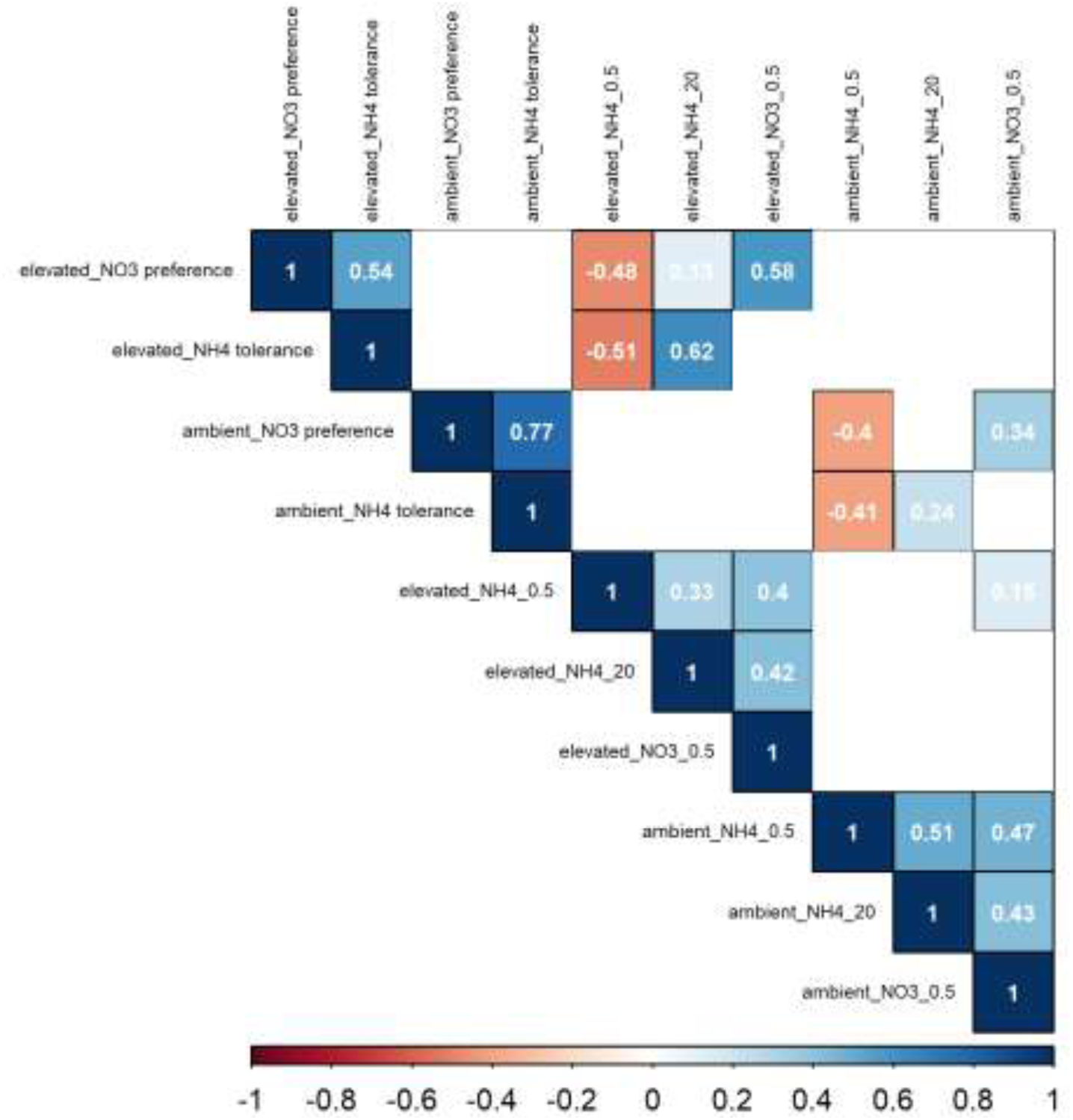
Correlation between biomass traits of 6 Nested Association Mapping populations by CO_2_ concentration. Pearson’s correlation coefficients at p-value < 0.01 between biomass traits by CO_2_ concentration. Insignificant correlations are not displayed. The horizontal bar displays reference color for negative correlations (red) to positive correlations (blue).

**Supplementary Figure 5.**
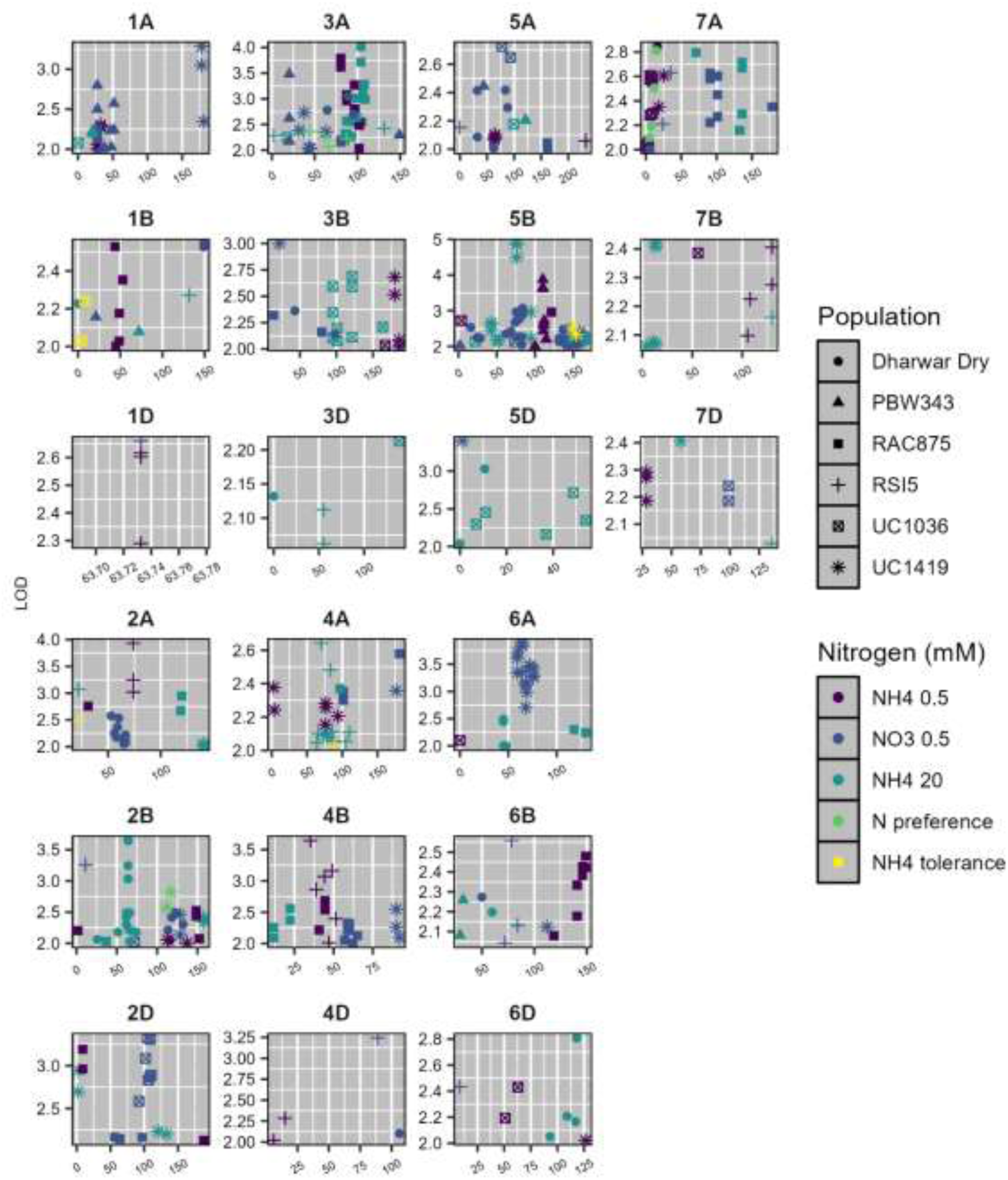
QTL mapping identified significant markers specific to populations and nitrogen forms across the genome. Distribution of significant markers (LOD > 2.0) across the wheat genome. Each panel represents a chromosome. Symbols represent individual bi-parental populations. Symbol colors represent nitrogen treatments or traits.

**Supplementary Figure 6.**
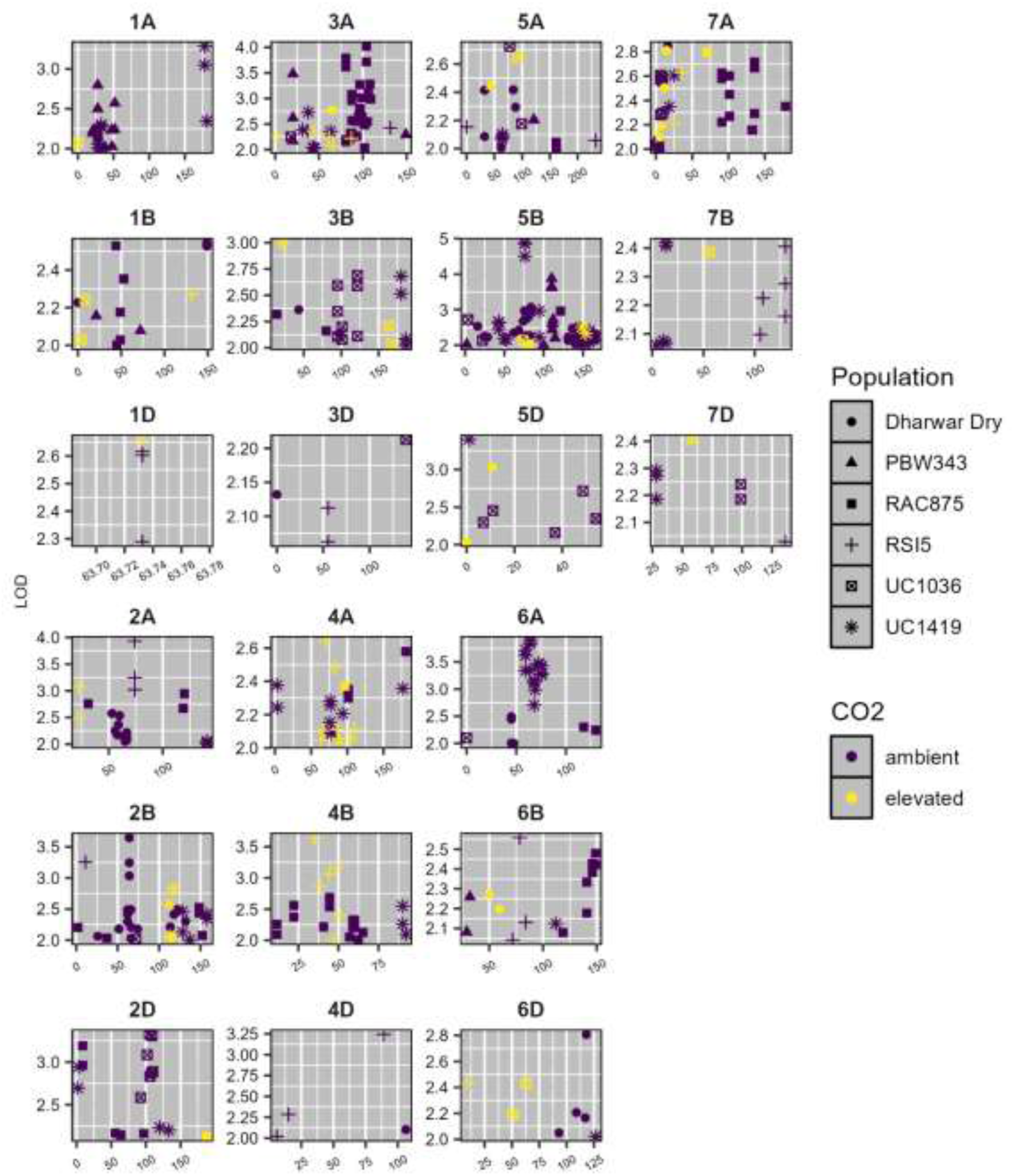
QTL mapping identified relatively more significant markers under ambient than elevated CO_2_ conditions. Distribution of significant markers (LOD = 2.0) across the wheat genome. Each panel represents a chromosome. Symbols represent individual bi-parental populations. Symbol colors represent CO_2_ levels.

---

**Supplementary Table 1.** Significant SNPs with p-value < 0.05. This table is available as a supplementary file.

**Supplementary Table 2.**
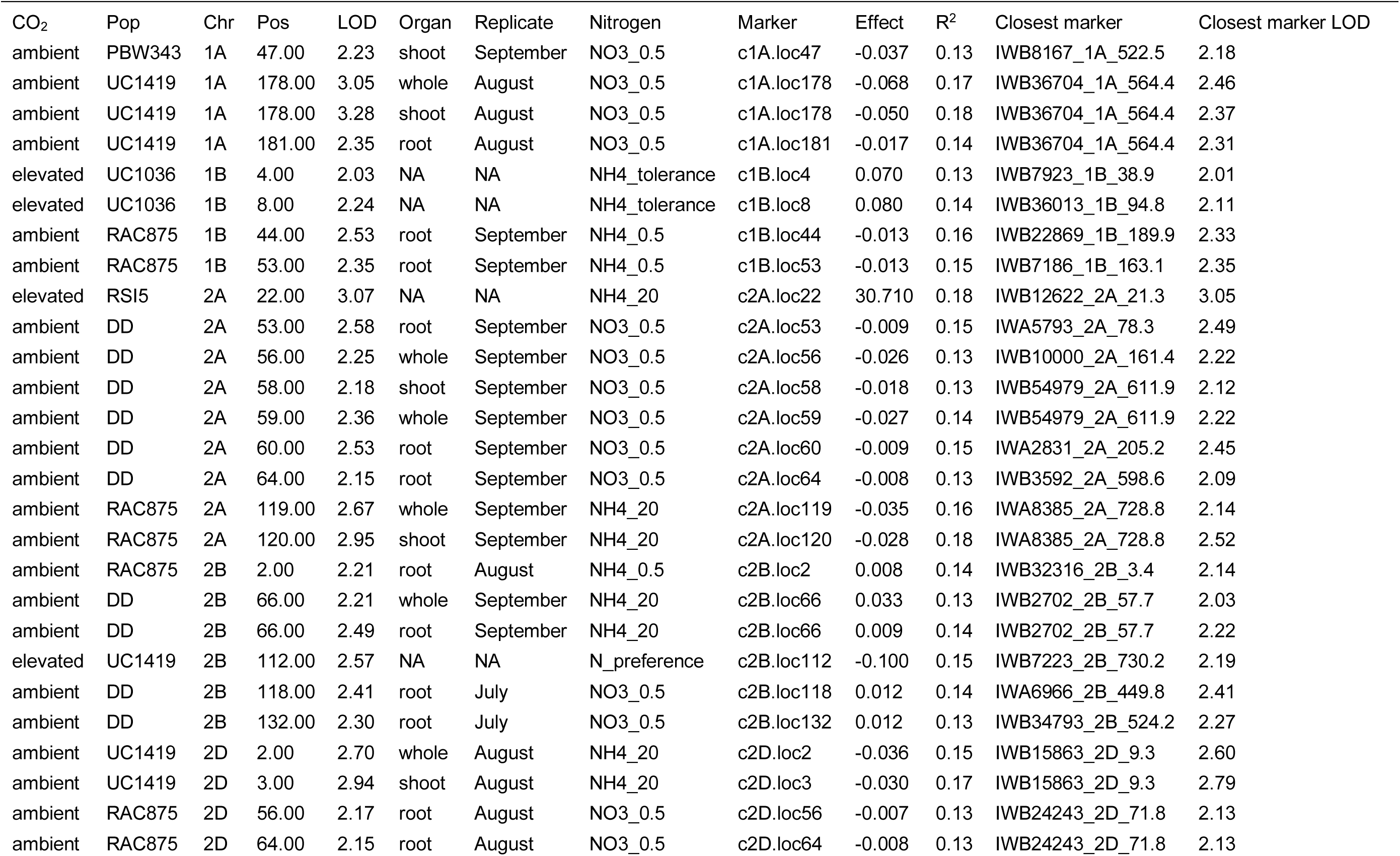
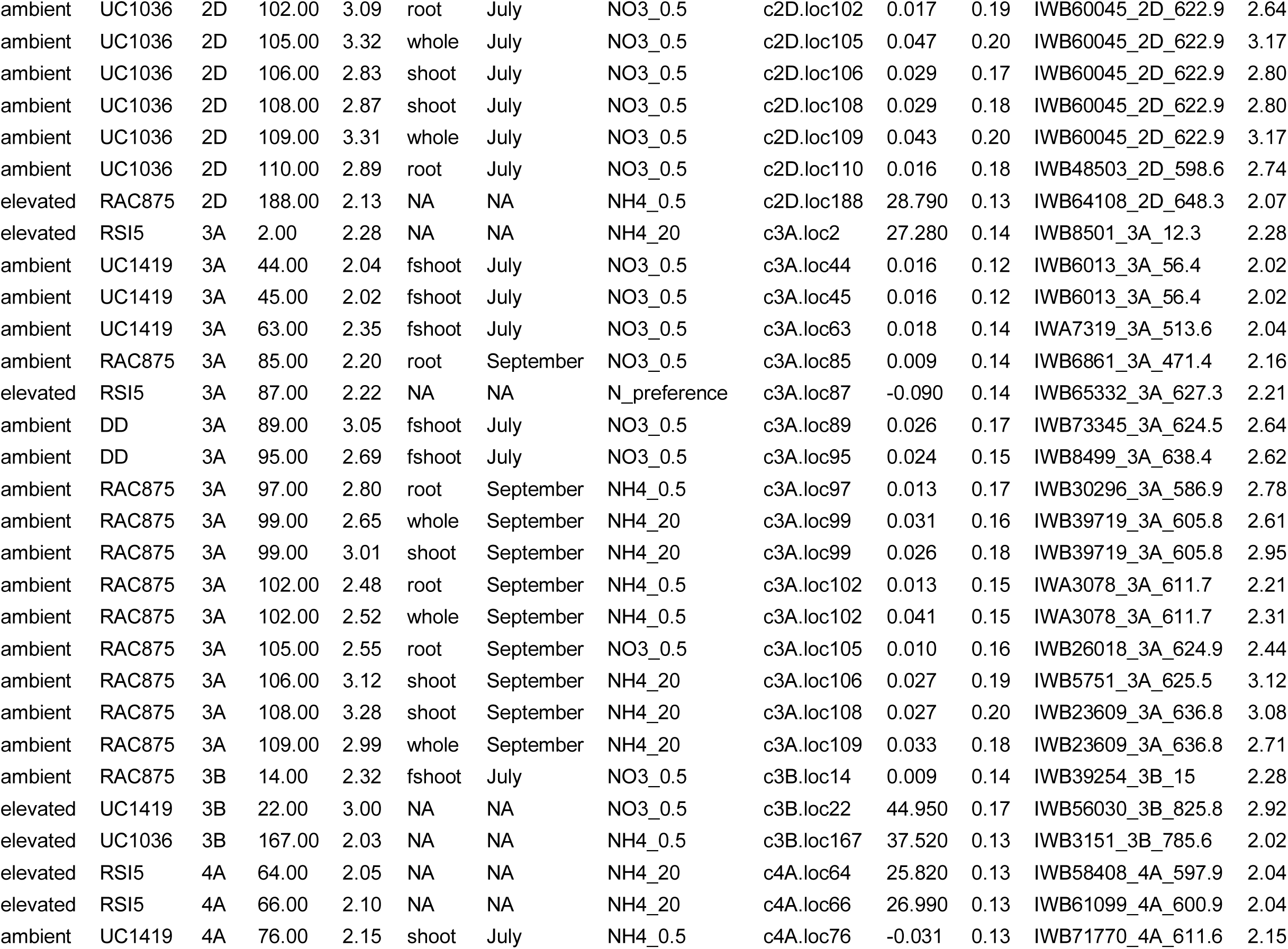
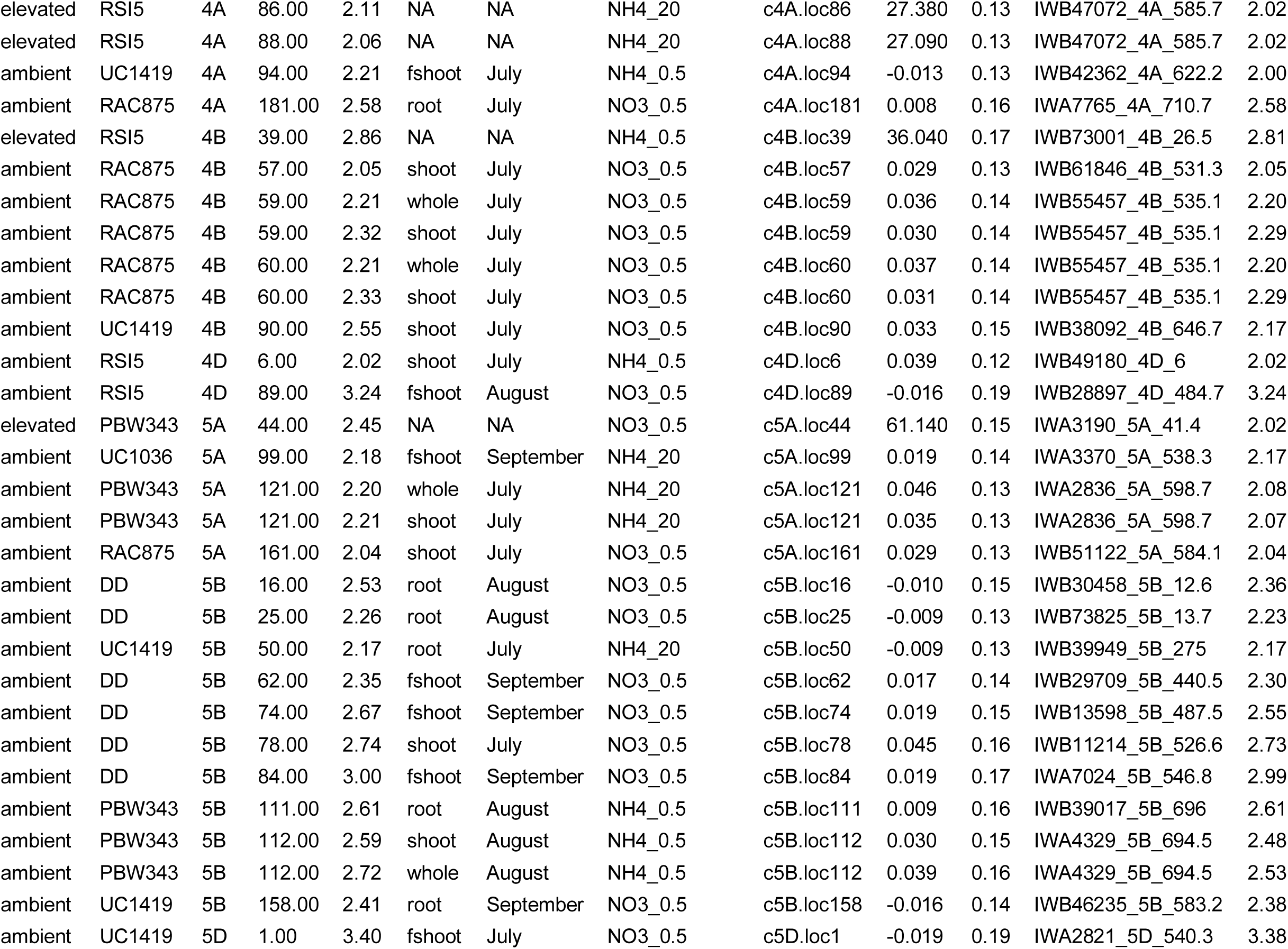
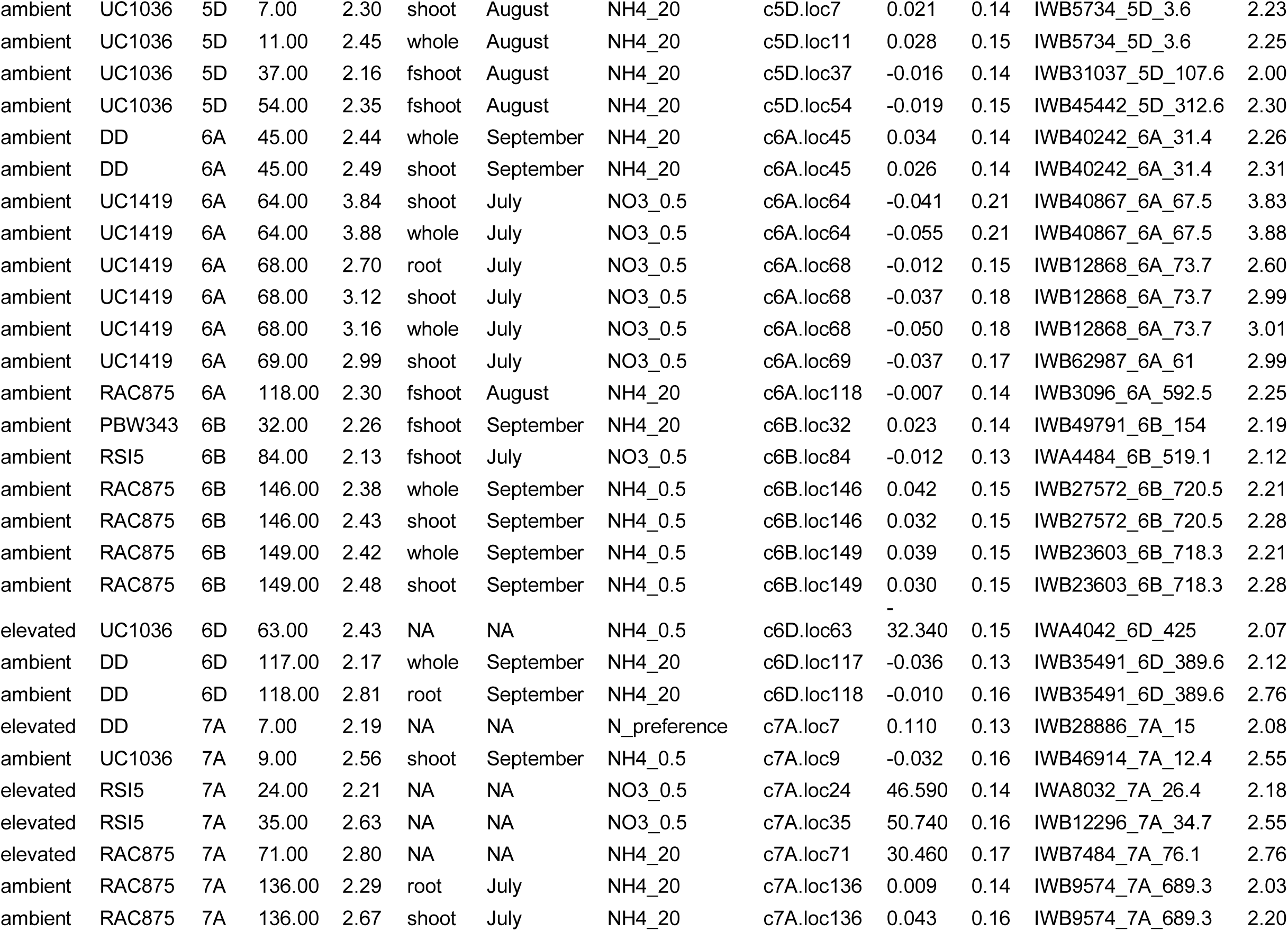
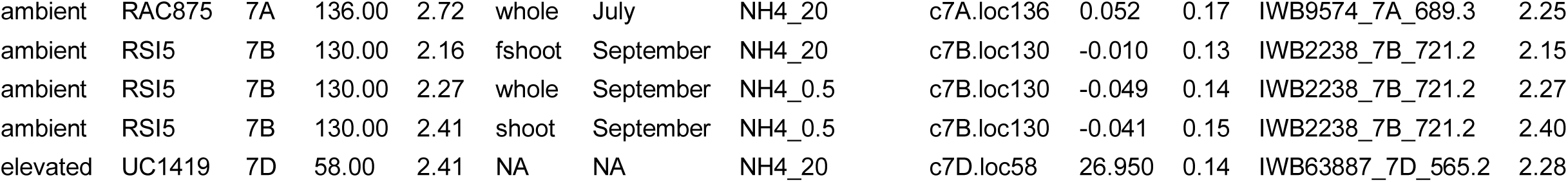
SNPs with LOD > 2.0 identified in QTL mapping of 6 biparental mapping populations at ambient and elevated CO_2_.

